# From apparent competition to facilitation, impacts of consumer niche construction on the coexistence and stability of consumer-resource communities

**DOI:** 10.1101/332437

**Authors:** Aurore Picot, Thibaud Monnin, Nicolas Loeuille

## Abstract

1. In addition to their direct trophic effects, some consumers have a positive indirect effect on their resource, due to niche construction. A predator can facilitate its prey resource acquisition, through prey transport, or through modifications of nutrient cycling. Other predators defend their prey against other predators, or actively manage it, as in agriculture, which is found in numerous taxa such as humans, but also ants, beetles, fishes and microbes.
2. Here we investigate the ecological consequences of considering such positive effects in a simple two resource–one predator module, in which the consumer has a positive effect on one of the resources.
3. We consider several scenarios, in which the positive effect of the resource is either non costly, ie resulting from a by-product of the consumer phenotype such as nutrient cycling, or costly. The cost either decreases the exploitation of the helped resource or the opportunity to forage the alternative resource.
4. We show that by modifying the trophic interactions in the module, niche construction alters the apparent competition between the resources, thereby impacting their coexistence.
5. We investigate how the intensity of niche construction impacts species coexistence, the distribution of biomass among the three species, and the stability of the community. When niche construction has little or no cost, it leads to higher consumer and helped resource densities, while decreasing the alternative resource density. Alternatively, when niche construction has a strong cost, the alternative resource can increase in density, niche construction thereby leading to the emergence of facilitative interactions among resource species.

## Introduction

While most studies in network ecology consider only one interaction type (ie, either food webs or mutualistic networks), the co-occurrence of different interaction types within networks has been of increasing interest in the last decade, in both theoretical and empirical studies (Fontaine et al., 2011; Kéfi et al., 2012; Mougi & Kondoh, 2012). In particular, positive interactions that co-occur with antagonistic interactions may originate from mutualism, facilitation (Bruno, Stachowicz, & Bertness, 2003), ecosystem engineering (Jones, Lawton, & Shachak, 1994) and niche construction (Odling-Smee, Laland, & Feldman, 1996). They can either directly affect the partner of interaction (as in mutualism) or alter the environment (as in ecological engineering) (Kéfi et al., 2012).

Importantly, the co-occurrence of multiple interaction types may affect predictions on general ecological questions such as ecosystem functioning, community stability and persistence (Fontaine et al., 2011; Kéfi et al., 2012). For instance, positive interactions can increase the persistence of a consumer under resource-limited conditions (Kylafis & Loreau, 2008) thus facilitating the coexistence and increasing community diversity (Gross, 2008). Positive interactions can notably occur when a given species has both trophic and non-trophic effects on the same partner of interaction. For instance, ant species that simultaneously tend aphids and prey on them can provide benefits to aphids by eating honeydew and protecting them from predators (Stadler & Dixon, 2005). Note that the frontier between a consumer helping its resource and a mutualist exploiting its partner is not clear, and the examples we cite later on could often belong to either categories (Bronstein, 2001; Offenberg, 2001). The net demographic effect of the interaction theoretically allows a complete classification but is not always measurable.

Indirect positive effects can emerge from a variety of consumer behaviours that improve some components of the resource growth rate (Brown, Ferris, Fu, & Plant, 2004 and references within). A first intuitive case would be a consumer facilitating its resource nutrient acquisition. For instance, while grazing, herbivores recycle nutrients to the soil, and may under some conditions increase primary productivity. An intermediate level of herbivory then leads to an optimal primary productivity (grazing optimization hypothesis, (de Mazancourt, Loreau, & Abbadie, 1998)). Predators may also reduce prey mortality, when they protect it against other predators (through interference with or predation of alternative predators) or inhibits the prey’s competitors. For instance, in devil’s gardens, ants kill competitors of their host plant species (Frederickson, Greene, & Gordon, 2005). Finally, a consumer may help its resource dispersal and reproduction (eg, seed dispersal linked to granivory, (Davidson, 1977)), or reduce prey intraspecific competition. The nematode *Caenorhabditis elegans* for instance transports its prey bacteria and inoculate them to unexploited resource pools (Ingham, Trofymow, Ingham, & Coleman, 1985; Thutupalli et al., 2017).

Such helping behaviours can occur at a cost for the consumer or not. This distinction ultimately changes the way the positive effect affects the demography of the consumer species in the system, altering the prey-consumer feedback. We call the positive effect passive if it results from a by-product of the consumer phenotype, with no direct metabolic cost. Nutrient cycling, as in the grazing optimization hypothesis (de Mazancourt et al., 1998), seed dispersal or passive transport of the resource to unexploited areas as in the nematode-bacteria interaction would all fall in this category (Thutupalli et al., 2017). Dissuading other predators from attacking the resource through the mere presence of the consumer could also be a passive positive effect.

When the positive effect on the resource is active, its negatively affects another consumer fitness component. For example, active resource management, that we refer loosely to as “agriculture”, incurs a cost in terms of time and energy. Agriculture - cultivation of plants, algae, fungi and animal herding - is found in humans, but also ants, beetles, fishes and even microbes (Boomsma, 2011; Hata & Kato, 2006; Mueller, Gerardo, Aanen, Six, & Schultz, 2005; Rowley-Conwy & Layton, 2011; Smith, 2016). The cost of agriculture can be envisioned through foraging theory (Charnov, 1976), where spending time on one resource reduces the available time to forage another resource, leading to an “opportunity cost”. Another type of cost can emerge if actively defending a resource against other predators or competitors implies moving away from this resource site or decreasing its consumption (eg, by allocating time to defense rather than consumption). Then, an “exploitation cost” scenario emerges with a trade-off between resource consumption and resource protection.

We here investigate the ecological consequences of considering such positive effects in a simple trophic module. We consider a consumer that feeds on a helped resource while also foraging on a second (non-helped) resource. Such simple trophic modules have been extensively used in ecology to understand mechanisms promoting coexistence and stability in ecological networks (Bascompte & Melián, 2005; Holt, 1997; Stouffer & Bascompte, 2010). We assume the consumer has a positive effect on one of the resources (the “helped” one). We consider three scenarios that cover passive positive effects (“no cost” scenario) and two types of active positive effects (“exploitation cost” and “opportunity cost” scenarios). “No cost” scenarios assume that niche construction only has an effect on the resource growth rate with no allocative cost (nutrient recycling, cross-feeding). In “exploitation cost” scenarios, we assume that investment in niche construction decreases the direct consumption of the helped resource. This may occur when time devoted to defense against predators constrains consumption (for instance, ants protecting aphids against ladybirds (Stadler & Dixon, 2005). “Opportunity cost” scenarios assume that niche construction decreases consumption of the alternative resource. This scenario is tightly linked to optimal foraging theory (Charnov, 1976; Pyke, Pulliam, & Charnov, 1977)(Charnov, 1976; Pyke et al., 1977) and exploitation-exploration trade-offs (Monk et al., 2018). It relates to transitions between predation and breeder behaviours, found in numerous species that specialize partly (facultative aphid rearing ants) or fully on cultivated resources (humans, obligate aphid rearing ants (Ivens, von Beeren, Blüthgen, & Kronauer, 2016), fungus growing ants (Chapela, Rehner, Schultz, & Mueller, 1994)).

We investigate how niche construction impacts species coexistence, the distribution of biomass among the three species, and the stability of the community. Predictions can be made considering the indirect effects occurring in the system. In our module, the two resource species do not directly compete but engage in apparent competition through their interactions with the shared predator (Holt, 1977). Any increase in biomass of a given prey has an indirect negative effect on the other prey as it increases predator density. Previous works show that the winner of the competition is the species that sustain the highest density of predator, leading to a *P\** rule similar to the *R\** of exploitative competition (Holt, Grover, & Tilman, 1994). Because we consider that one species receives an additional positive effect from the consumer, indirect effects are altered. If the net effect of the consumer on the helped resource is positive, then the apparent competition may become an apparent antagonism as the alternative resource has a positive indirect effect on the helped resource (see figure 1). In “no cost” scenarios, we predict that niche construction increases the growth rate of the helped resource, hence negatively impacting the alternative resource through increased apparent competition. Eventually, such an effect may lead to the loss of coexistence. Considering a trade-off with the consumption of either resource may modify these predictions by affecting the balance of indirect effects. In the “exploitation cost” scenario, niche construction decreases the consumption of the helped resource, hence makes it less vulnerable to predation: we predict that it would win the competition because it suffers less from the indirect negative effects received from the alternative resource. We therefore predict that niche construction should eventually negatively impact coexistence. In the “opportunity cost” scenario, niche construction decreases the consumption of the alternative resource, hence the effects of apparent competition should be more balanced among prey species, promoting coexistence.

**Figure 1:**
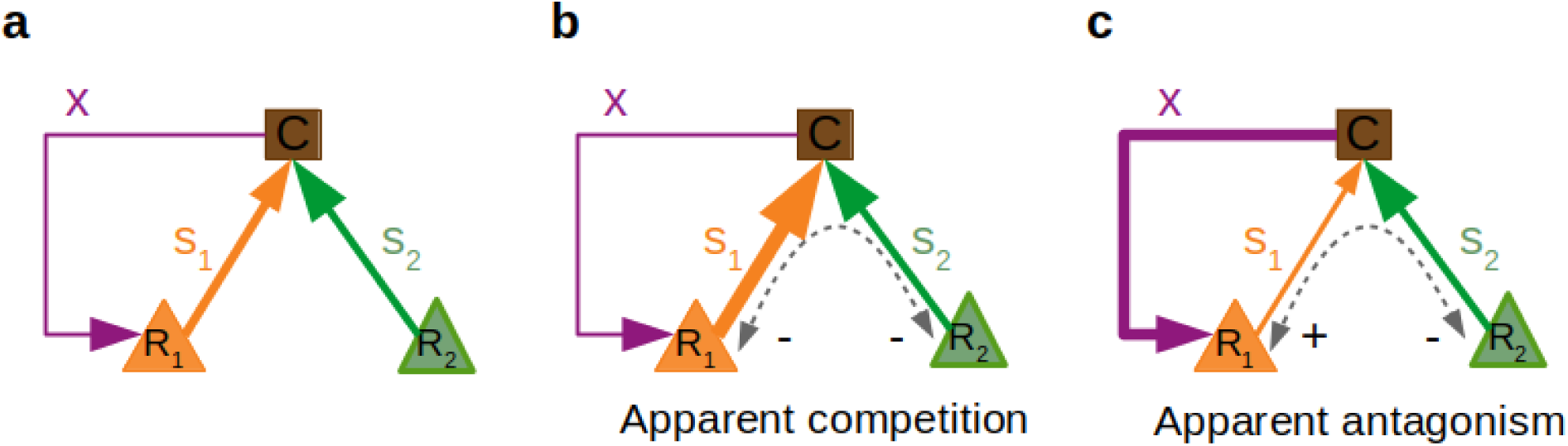
Presentation of the three-species model and indirect effects occurring between the resources. **a** Direct interactions: the consumer C consumes both resources R_1_ and R_2_ depending on its specialization on each resource (s_1_ and s_2_). It increases the growth of resource R_1_ by a factor x, which is the niche construction investment trait. **b** When the outcome of the C-R_1_ interaction is negative (s_1_>x), resources are limited by apparent competition. **c** When the outcome of the C-R_1_ interaction is positive (s_1_<x), the resources engage in apparent antagonism.

Predictions can also be made regarding the effects of niche construction on the stability of the system. In the “exploitation cost” scenario, niche construction should reduce the energy flux from the resource to the consumer, relative to the consumer loss. Such limited energy fluxes should be stabilizing, as long as the net interaction remains trophic (Rip & McCann, 2011). Considering the whole three-species module, niche construction is expected to be stabilizing when it increases interaction heterogeneity among prey species (McCann, Hastings, & Huxel, 1998), for instance due to costs on the consumption of either species. However, if the net interaction between the consumer and the helped resource becomes mutualistic due to large positive effects, we expect it to be destabilizing, because a negative trophic feedback loop then becomes a positive feedback loop (May, 1973).

## Model presentation

Ecological dynamics are modeled using ordinary differential equations (eq 1):

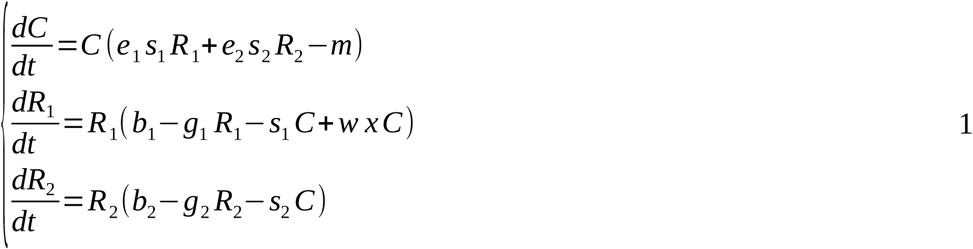

The consumer *C per capita* growth rate depends on its consumption of resources (modelled by specialization on resource *R*_*i*_, *s*_*i*_, and conversion efficiency of resource *R*_*i*_, *e*_*i*_) and on its *per capita* death rate *m*. Resources have a logistic growth in the absence of the consumer (allowing for their coexistence in such situations): *b*_*i*_ is the *per capita* birth rate and *g*_*i*_ is the intraspecific competition rate for resource *R*_*i*_. We do not consider any direct competition between resources so that their coexistence can be entirely understood based on variations of indirect effects inherent to the consumer species. To this logistic growth is added a consumption rate scaled by the specialization of consumer on the resource, and a niche construction effect for resource *R*_*1*_. All interactions are linear. In particular, in this first model, niche construction is proportional to the investment trait *x*, and the consumer density, modulated by a niche construction efficiency *w*. Such simple linear functions allow for an analytical study of the system. In the Supplementary Information, we consider a saturating response for the niche construction effects and discuss the robustness of the results we present in the main text. We assume that *e*_*i*_, *s*_*i*_, *b*_*i*_, *g*_*i*_, *x* are positive, so that niche construction is facultative for the maintenance of resource 1.

The ecological analysis of the system focuses on the feasibility and linear stability criteria applied on the different possible equilibria. Feasibility conditions require the positivity of all equilibrium densities. The stability analysis relies on the analysis of the Jacobian matrix (thereby assessing the return time to equilibrium following a small disturbance) and of the invasibility of considered equilibrium by species that are not present at equilibrium. All figures and computations were made using Mathematica 11.

## Results

The model displays community states (equilibria) at which resources can subsist without the consumer, or the consumer coexists with one or both resources. We sum up the general stability and feasibility conditions for the “no cost” scenario, and study how niche construction impacts these coexistence and stability conditions. We then investigate how the addition of a cost modifies those results in the “exploitation cost” and “opportunity cost” scenarios. The detailed mathematical analysis can be found in the Supplementary Information.

### 1) Coexistence and stability under the “no cost” scenario (*s*_*1*_*’(x)=s*_*2*_*’(x)=*0)

In this scenario, the positive effect of the consumer on its resource happens through a passive effect, as a by-product of metabolism or activity of the consumer. There is no direct cost for the consumer. For instance, the large effects on nutrient recycling by the wildebeest in the Serengeti ecosystem (McNaughton, 1976) could be considered as a motivation for such a scenario. The positive effect only impacts the growth rate of resource 1 through the consumer density-dependent factor, and predation rates *s*_*1*_ and *s*_*2*_ are constant.

We show that the consumer-helped resource (*C-R*_*1*_) equilibrium is feasible and stable when the interaction between the two species remains trophic, (ie, positive effects are not too high). The ratio between the resource birth rates and their vulnerabilities determines the invasion potential for resource *R*_*2*_: if it has a high birth to vulnerability ratio, it can invade the consumer-helped resource system. In this (*C-R*_*1*_) subsystem, niche construction has no effect on the resource density but increases the consumer density. The increase in growth rate is compensated by an increase in predation rate, so that the resource remains top-down controlled.

Regarding the (*C-R*_*2*_) equilibrium, our analysis reveals that niche construction does not have an impact on the equilibrium densities, as the helped resource is absent. Niche construction is destabilizing, as large *x* allows an invasion of the equilibrium by resource *R*_*1*_. Conditions of coexistence of the three species can therefore be expressed as upper and lower limits of the positive effect intensity. Hence, niche construction favors coexistence at intermediate values.

Niche construction affects the distribution of species densities. Given the dynamical system 1, species densities at the coexistence equilibrium in the “no cost” scenario can be written:

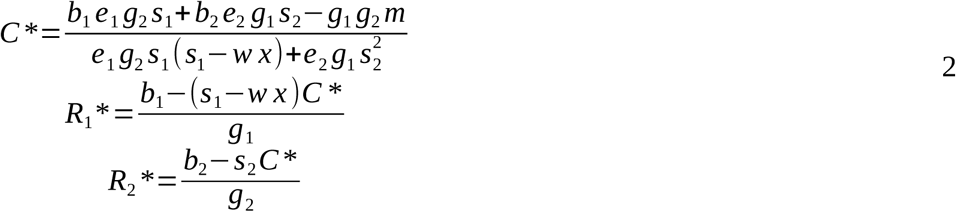

It is then possible to show how species densities vary depending on the intensity of niche construction:

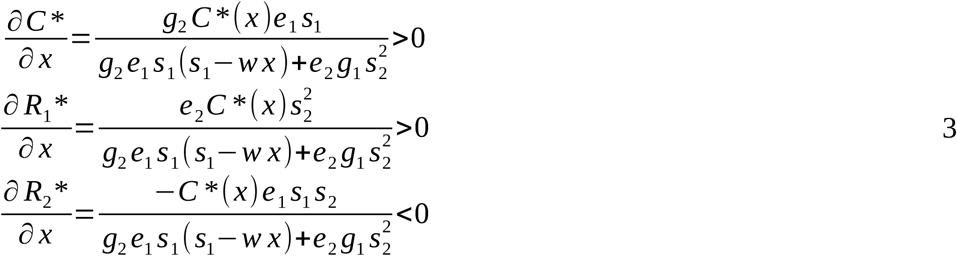

As predicted, niche construction has a positive effect on the helped resource density, leading to a positive bottom-up effect on its consumer, negatively affecting the alternative resource density through apparent competition (figure 2). Equilibrium densities at the *C-R*_*2*_ equilibrium do not vary with the intensity of niche construction, but the invasion potential of *R*_*1*_ increases up until *R*_*1*_ can invade, eventually leading to coexistence.

**Figure 2:**
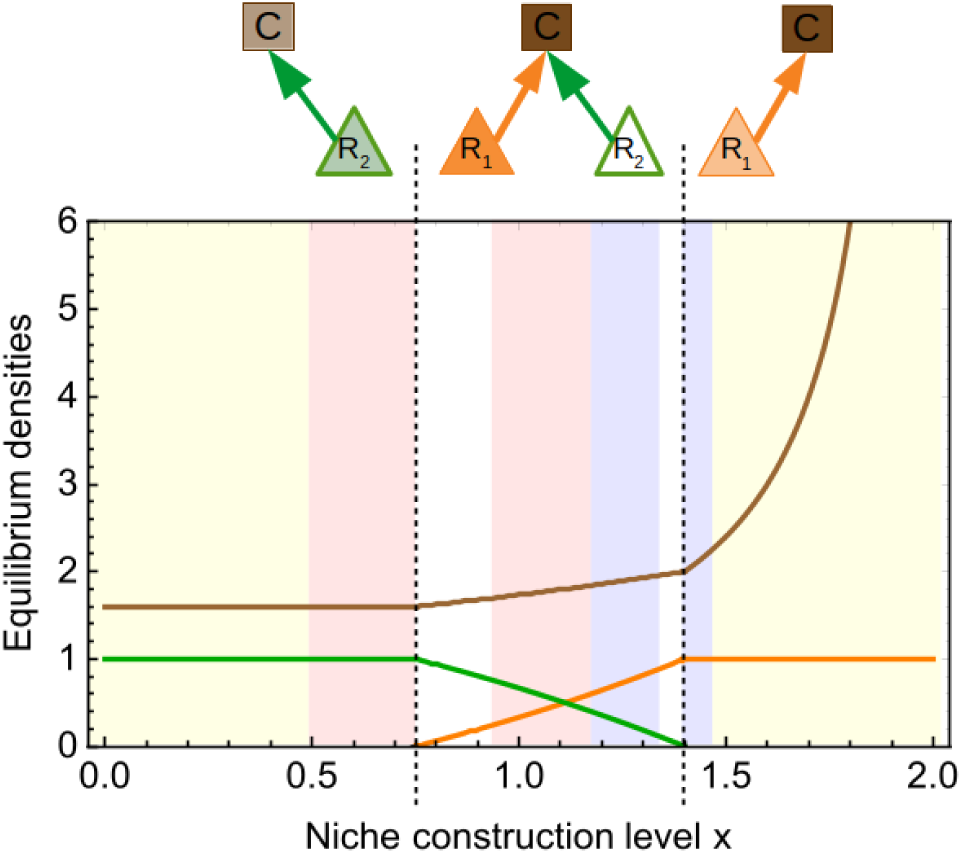
Effects of niche construction on the stable equilibrium densities and stability in the “no cost” scenario. The consumer density is in brown, resource 1 density in orange, resource 2 density in green. Above the plot, the different states of the module are represented. An empty box means that density decreases, light box means that density does not vary, while a dark box means that density increases. Within the plot, the background shows stability variation: no variation (yellow), destabilization (red), stabilization (blue). Stability is non-monotonous in white areas (transition between ecological states). e_1_**=**e_2_=0.5, g_1_=g_2_=0.8, m=1, b_1_=2, b_2_=4, s_1_=2, s_2_=2.

Niche construction also affect the resilience of the system (measured as the negative real part of the dominant eigenvalue (Pimm & Lawton, 1977)). The stability measure varies abruptly around the ecological states frontiers (see Rip & McCann, (2011)), and we focus on variation for intermediate densities. For the coexistent equilibrium (as predicted from the mathematical conditions, see (S12)), niche construction is initially destabilizing then stabilizing, as it eventually increases interaction heterogeneity (McCann et al., 1998). Concerning the *C-R*_*1*_ equilibrium, niche construction is stabilizing, as it decreases the *per capita* energy flux from the resource to the consumer (Rip & McCann, 2011).

### 2) Effects of costly niche construction on coexistence and stability

We now assume that niche construction is costly for the consumer. This cost may decrease the consumption of either the helped resource (“exploitation cost”), or the alternative resource (“opportunity cost”). “Exploitation cost” scenarios include situations where higher positive effects on the helped resource (eg agriculture) decreases the exploitation of the same resource. This can happen because the time or energy devoted to protection of the resource against predators cannot be used for consumption (for instance, ants protecting aphids against ladybirds (Stadler and Dixon 2005)). “Opportunity cost” scenarios correspond to situations where the foraging on an alternative resource decreases, implying a trade-off between predation and agriculture activities.

From eq 1, it is easy to show that niche construction affects the distribution of species densities as (see Supplementary Information (S14)):

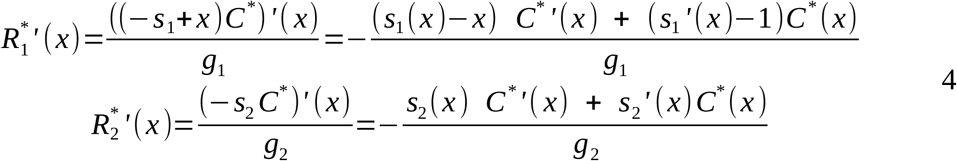

These derivatives are made of two terms. The first one shows the ecological consequences of niche construction (its impact on consumer population *C*’(x)*), while the second term embodies the effect of each cost, *s*_*1*_*’(x)* and *s*_*2*_*’(x)*.

#### a) “Exploitation cost” scenario (s_*1*_*’(x) < 0, s*_*2*_*’(x) = 0)*

In the “exploitation cost” scenario, niche construction directly lowers the consumption of *R*_*1*_. Intuitively, such a decrease in consumption reinforces the positive effect of niche construction on *R*_*1*_. In turn, this should increase apparent competition and harm *R*_*2*_. In such a scenario, we obtain that:

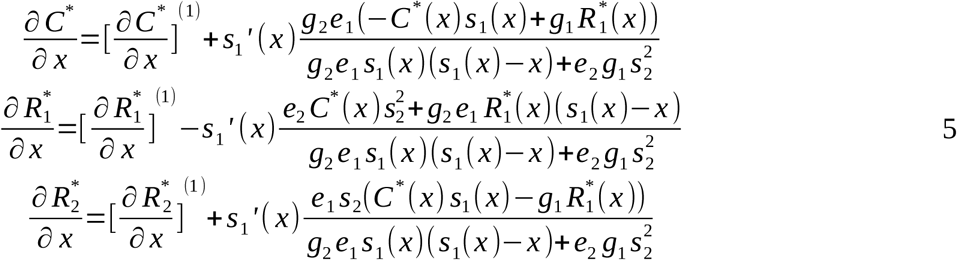

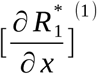, 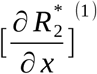 and 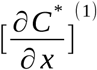 are density variations in the “no cost” scenario (see eq. 3). These derivatives can either be all positive or all negative. Because niche construction comes at a cost for the consumer, the overall effect on *C\** may be negative. Consumption is then relaxed on the alternative resource so that it may also benefit from niche construction.

On figure 3, we illustrate these outcomes assuming a linear trade-off: *s*_*1*_*(x) = s*_*1*_ *– β*_*1*_ *x,* with different trade-off strengths (*β*_*1*_) and interaction intensity *s*_*2*_. At low niche construction, when the alternative interaction is low (panels a and b), variations are similar to the no cost scenario (equation 3 and 5). On panel a, note that intermediate niche construction does not allow the stable maintenance of a coexistent system (grey area). Starting from a coexistent state, dynamics lead to the extinction of resource *R*_*2*_ and the exponential growth of the *C-R*_*1*_ system. An initial *C-R*_*1*_ subsystem (*R*_*2*_=0) would lead to the extinction of the consumer. This may be explained by the fact that in the first case, the transient dynamics during which *R*_*2*_ is present allows large consumer populations that ever increase due to the positive feedback with *R*_*1*_ (the interaction *C-R*_*1*_ is mutualistic, as *s*_*1*_*<x* in this area). In the second case, the initial state does not allow such an infinite growth as populations are too small. Because *x* is intermediate, the cost on the consumer growth rate is not high enough to stabilize the dynamics: the energy intake from the consumption of resource 1 is still high. Hence, we obtain typical unstable mutualistic dynamics with exponential growth of the consumer and the helped resource (those dynamics are illustrated in supplementary information, figure S1). When niche construction *x* is high (fig 3a, right), it decreases consumer density (reduced energy intake). Since the interaction is mutualistic, this also decreases *R*_*1*_ density. From the helped resource perspective, the positive effect of niche construction (increasing *x* and reducing attack rate *s*_*1*_) is compensated by the negative density-dependent effect on the consumer. Considering equation 4 helps visualizing this balance: the first part of the equation is negative because *C*’(x)* < 0 and *s*_*1*_*(x) – x* < 0 (the interaction is mutualistic). The second part corresponds to a positive effect stemming from the reduction in attack rate *s*_*1*_*(x).* On the contrary, variations in *R*_*2*_ density is only driven by the consumer density effect (equation 4). We would like to draw the attention on an important consideration here: for high values of niche construction, the *C-R*_*1*_ interaction is mutualistic but it is not symmetrical. In particular, when *x* is high enough, *s*_*1*_*(x)* tends towards 0 so that the interaction tends to commensalism (null effect of *R*_*1*_ on *C* while *R*_*1*_ benefits from niche construction). In panel c, because the alternative interaction is high, *R*_*1*_ dominates the apparent competition when there is low niche construction. Increasing *x* first increases both *R*_*1*_ and *C* but eventually heightens the cost of niche construction. *R*_*2*_ can eventually invade the system when consumer density goes below its *P*.* Niche construction then favors both resources densities while harming the consumer density.

**Figure 3:**
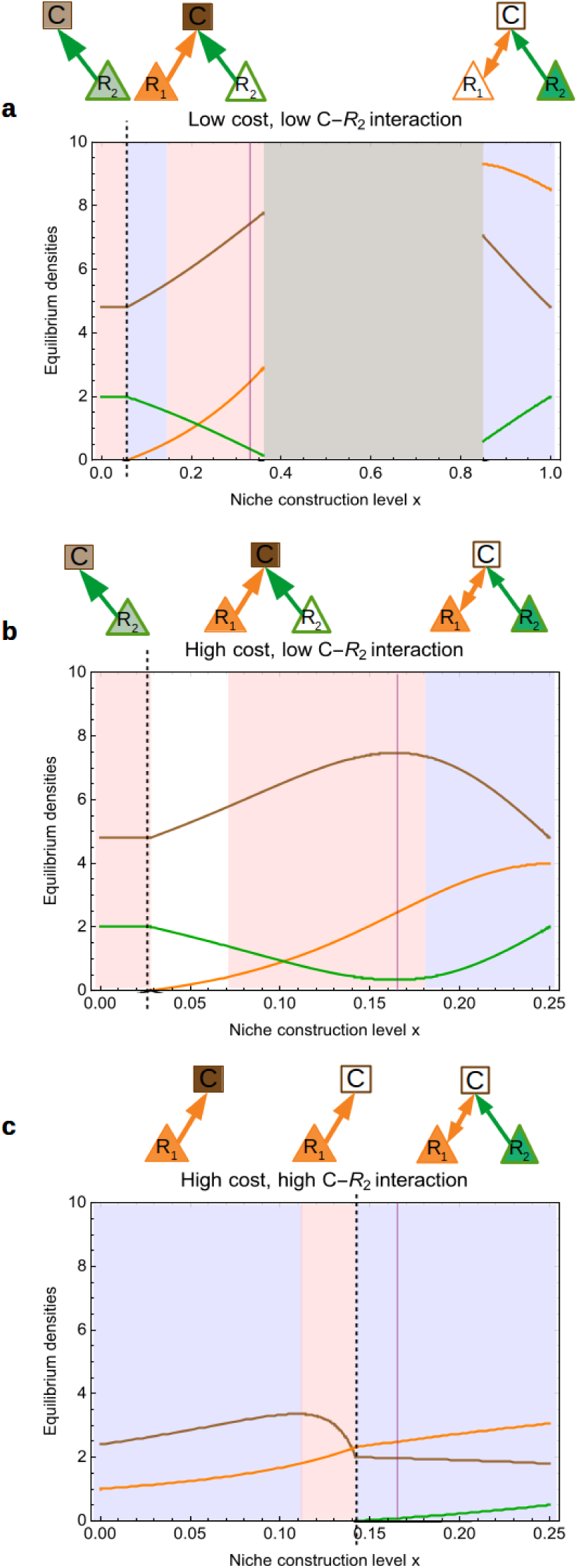
Effect of niche construction in the “exploitation cost” scenario for cost values and alternative resource interactions. The color code is the same as for figure 1. The grey area corresponds to no stable equilibria. The purple line indicates the value of niche construction where interaction between C and R_1_ switches from trophic to mutualistic. e_1_**=**e_2_=1, g_1_=g_2_=0.8, m=1, b_1_=2, b_2_=4. In panel a, s_1_(x)= 0.5 – 0.5 x, s_2_ = 0.5. In b, s_1_(x)= 0.5 – x, s_2_ = 0.5. In c, s_1_(x)= 0.5 – x, s_2_ = 2.

Effects of niche construction on stability are largely consistent with the “no cost” scenario (compare Fig3a,b *vs* Fig2). However, note that for higher *x* values, niche construction is stabilizing. Niche construction costs there reduce the attack rate on *R*_*1*_ and makes the interaction tend to commensalism. This reduces the positive feedback between *R*_*1*_ and *C* thereby explaining the stabilization of the system. As a corollary, intermediate niche construction levels lead to unstable dynamics such as in the grey area of Fig3a.

#### b) “Opportunity cost” scenario (s_*2*_’(x) < 0, s_*1*_*’(x) = 0)*

Given “opportunity costs”, niche construction decreases predation on the alternative resource. We can expect this to dampen the effects of apparent competition: as *x* increases, benefits on resource *R*_*1*_ increase, and predation on *R*_*2*_ is relaxed. We thus expect coexistence to be facilitated in this scenario compared to the previous ones.

We obtain that:

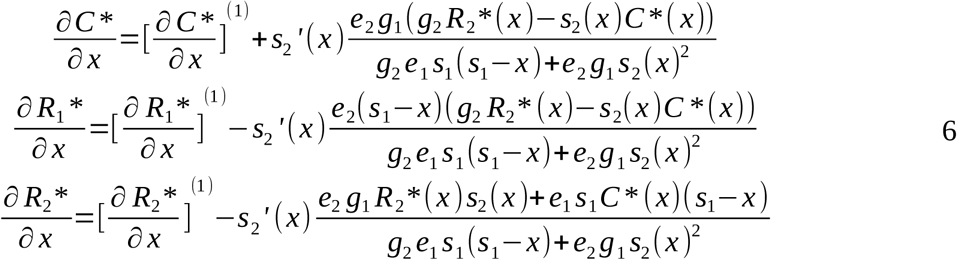

As in the “exploitation scenario”, we illustrate the effects of the cost, assuming a linear trade-off *s*_*2*_*(x) = s*_*2*_ *– β*_*2*_ *x* (Fig4), varying the intensity of the cost and interaction *s*_*1*_. From system 6, *s*_*2*_*’(x)* modifies the density variations expected from the “no cost scenario”. When this cost is low (*β*_*2*_∼0), qualitative patterns are consistent with the no cost scenario, both in terms of density and stability variations (Fig4a). Higher costs benefit all densities, as predicted (Fig4b). Niche construction also stabilizes the system, as it increases the asymmetry between the two interactions (Fig4b,c). Results in Fig4c, where both the cost and the alternative interaction are high, are consistent with those of figure 3c, that also assumed a high cost and a high alternative interaction. *R*_*1*_ can only invade when high niche construction reduces the consumer density under its *P\**. The two resource densities then increase with *x*, while consumer density slightly decreases. At low *x* values, for the consumer-*R*_*2*_ equilibrium, niche construction is first stabilizing then destabilizing (eventually allowing the invasion of *R*_*1*_). Results are also similar at higher levels of niche construction (fig 4c vs 3c).

**Figure 4:**
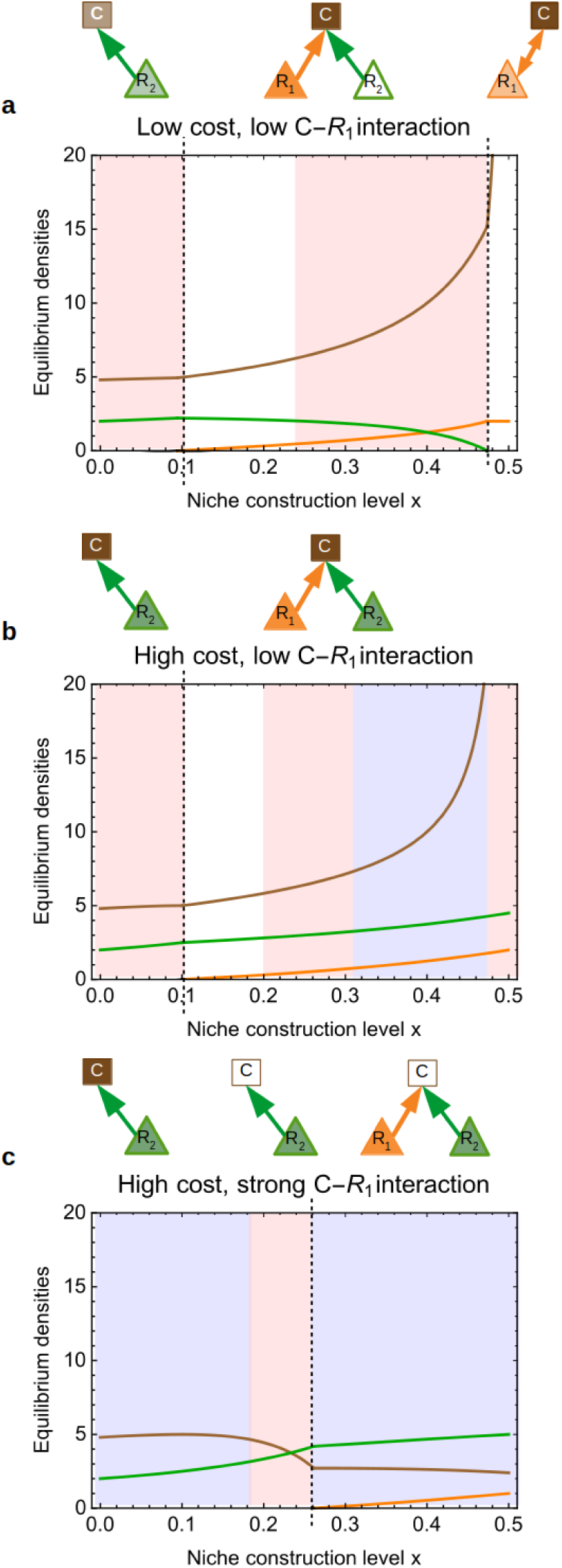
Effects of niche construction in the “opportunity cost” scenario. The color code is the same as in figure 3. e_1_**=**e_2_=1, g_1_=g_2_=0.8, m=1, b_1_=2, b_2_=4. Panel a: s_2_(x) = 0.5 – 0.5 x, s_1_ = 0.5, panel b: s_2_(x) = 0.5 - x, s_1_ = 0.5,, panel c: s_2_(x) = 0.5 – x, s_1_ = 1.

The general results for the stock variations of the densities with *x* in the three scenarios are summed up in the Table 1. We note that, when variations can be determined, the consumer is usually positively affected by niche construction, except in high cost scenarios. Similarly, niche construction is often positive for the helped resource, and may create positive facilitative effects on the second resource, when costs of niche construction are included.

**Table 1:**
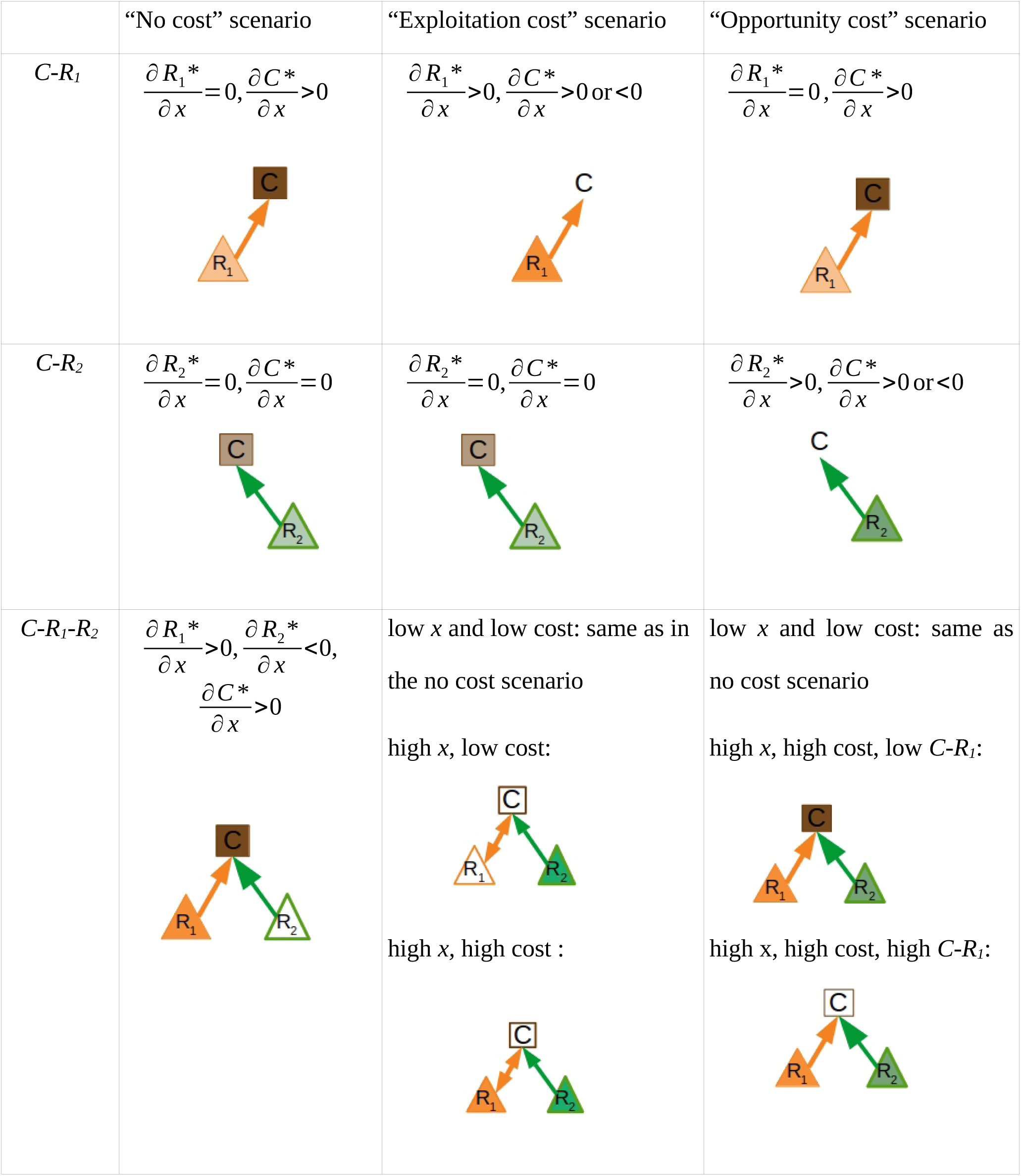
General effects of niche construction on density distributions. Empty, dark, light and no boxes mean decrease, increase, no variation and indetermination, respectively.

### 3) Robustness analysis

While we assumed effects of niche construction to be proportional to average investment *x* and consumer density, little information exists to link niche construction with species demography. In the supplementary material, we study an alternative model where niche construction eventually saturates. Effects of niche construction on the distribution of biomasses are largely consistent with the linear model, though species density variations are more moderate and stabilize because of the saturating response.

## Discussion

The motivation for our study was to investigate how the addition of a non trophic positive effect to a trophic interaction impacts the structure and functioning of a one-consumer-two-resources module. We investigate three scenarios that differ in whether and how the positive effect entails an allocative cost for the consumer. We discuss the three questions that we addressed in the introduction: the effect of niche construction on coexistence, on the distribution of densities within the module, and on stability.

When niche construction has no or little cost, we find that niche construction generally favors coexistence when the alternative resource initially dominates the apparent competition (for instance because it has a higher intrinsic growth rate or suffers less predation) (Holt, 1977). Niche construction then benefits the helped resource and allows coexistence, though increasing further eventually leads to the competitive exclusion of the alternative resource. Such a pattern may be linked to various empirical examples. For instance, devil’s gardens are almost pure plantations of one tree species maintained by ants killing other, more competitive plants (Frederickson et al., 2005). On the contrary, if the two resources are initially equally competitive or the alternative resource is less competitive, niche construction increases the asymmetry between the two resources thereby limiting coexistence. These results are consistent with classic trophic module studies (Holt & Lawton, 1994), with niche construction simply modulating apparent competition.

Niche construction also alters the abundance distribution of the different species. Intuitively, we expect that niche construction behaviour may benefit the consumer population as well as the abundance of the helped species. When niche construction has no or little cost, we indeed find such a pattern, and also find that the alternative resource density is decreased. Such results are largely consistent with the direct effects of niche construction: it has a positive effect on the helped resource density, while increasing consumer density through bottom-up effects (increased resource availability). This increase in consumer biomass leads to a negative top-down effect on the alternative resource density. Consistent with such a mechanism, some studies where ant tend aphids have noted an increased predation from ants on alternative non-tended aphid species (Warrington & Whittaker, 1985). It is however not clear whether this negative effect occurs from a consumer density increase or through changes in the foraging pattern. Similarly, Wimp & Whitham (2001) show that the experimental removal of an aggressive aphids-tending ant strongly increases the biodiversity of other arthropod species, suggesting that the ant-aphid association has a negative impact on the abundance of other species and may indeed limit species abundances and coexistence.

Interestingly, larger costs can strongly modify this intuitive pattern. The main effect of a strong cost is, counter-intuitively, not dependent on where this cost occurs in our trophic module. Whether this cost involves the exploitation of the helped resource (strong “exploitation costs”), or of the alternative resource (strong “opportunity costs”), it leads to a negative effect of niche construction on consumer density, and a positive effect on the alternative resource. The effect on the helped resource varies depending on the balance between the direct niche construction effect (through trait *x*) and the indirect effect on consumer density. This general pattern can be interpreted in the following way: if the interaction between the consumer and the helped resource becomes globally mutualistic (ie, the positive effect is larger than the trophic effect of the consumer), then the consumer and helped resource densities covary with the intensity of niche construction. Given a high cost, niche construction then decreases both densities, helping the alternative resource through relaxed predation (either because of the decrease in consumer density or, in the case of opportunity costs, because of lower predation rates). If the interaction remains mainly trophic, then niche construction generally decreases the consumer density but increases both of the resource densities due to relaxed predation. Such overall positive effects of niche construction on all resources can be linked to the notion of prudent predation (Goodnight et al., 2008; Slobodkin, 1974) We also note, in the “exploitation cost” scenarios, that consumer density reaches an optimum at intermediate niche construction.

We note that such positives effect on the alternative resource, not directly helped, may be seen as a facilitation of this species by the *C-R*_*1*_ interaction. Such a facilitation emerges when the interaction between the consumer and the helped resources becomes mostly mutualistic. Assuming strong costs of niche construction, apparent competition is then replaced by a dominant facilitation effect. Assuming strong costs, niche construction leads to an increase in both resources because it relaxes predation impacts not only on the helped resource, but also on the alternative resource species (either through a consumer density-dependent effect, or through a decrease in the *per capita* attack rate). Our model thus allows a continuum between competitive and facilitative interactions, whose importance in ecology is increasingly recognized (Bruno et al., 2003; He, Bertness, & Altieri, 2013; Kéfi et al., 2012). Because the balance between competition and facilitation in our system depends on the levels of cost, this highlights the importance of investigating the possible trade-offs associated to niche construction. We stress that the non-cultivated resource is facilitated though it receives no direct benefit. Niche construction can modify a foraging pattern in a way that is not necessarily costly: for instance, fire ants forage on the ground but when they tend aphids they also forage on the arthropods present on the host plant (Kaplan & Eubanks, 2005). Foraging on plants for preys only is not profitable but foraging for honeydew makes foraging for nearby preys profitable. We here assume that costs of niche construction act on predation rates. This assumption, based on allocation of time and energy between different functions (niche construction on one side, foraging on the other), could be generalized, for instance assuming a continuum between our “exploitation cost” and “opportunity cost” scenarios.

Effects of niche construction on stability are less intuitive. When costs are low, niche construction is first destabilizing then stabilizing. This can be explained thanks to previous works on stability in trophic modules or networks. Notably, when it increases the heterogeneity of interaction strengths (eg, due to costs), niche construction is stabilizing, as found in food web modules (McCann et al., 1998). When only the helped resource is present, increasing niche construction is also stabilizing, because it decreases the *per capita* energy flux from the resource to the consumer, consistent with classical “paradox of enrichment” results (Rip & McCann, 2011; Rosenzweig, 1971).

We chose to keep our model linear to allow for a better analytical tractability. We assume no direct competition among resources focusing on apparent competition instead. We believe that adding direct (or exploitative) competition between the resources would give predictions similar to the *P*-R\** coexistence rule (Holt et al., 1994) with niche construction modulating the *P\** value of the resource species. We analyze the robustness of the model regarding the assumption of a linear niche construction effect, investigating a scenario where effects of niche construction progressively saturate. In terms of equilibrium densities response, we find similar patterns in the linear and the saturating model. The saturating response is generally stabilizing, as expected from previous studies showing the stabilization of mutualism when considering saturating functional responses (Holland & DeAngelis, 2010; Holland, DeAngelis, & Bronstein, 2002).

The explicit consideration of trophic interaction modifications has recently received increased attention (Terry, Morris, & Bonsall, 2017 and references within). In this line of work, our model investigates how positive niche construction effects interfere with apparent competition, to constrain species coexistence and community stability. While these small modules are by essence simplified compared to larger, natural networks, ecology has a long tradition of using them to propose and test predictions on coexistence or stability. We also assume that the consumer positively affected one of the two resource species only. Note however that a consumer often has positive effects on several of its resources, a case we can reasonably expect in the cases of nutrient cycling (Cargill & Jefferies, 1984; de Mazancourt et al., 1998) and seed dispersal (Serrano-Cinca, Fuertes-Callén, & Mar-Molinero, 2005; Thutupalli et al., 2017). Explicit simulations of these more complex (ie, more species or more diffuse effects) scenarios are beyond the scope of this article, but they would help to get a better understanding of the role of non trophic interactions in larger networks, by allowing the accumulation of more indirect effects of different types (Kéfi et al., 2012).

While the present study would benefit from being extended to multispecies networks, we believe that even in such situations, some of our results would hold. For instance, if a herbivore has a positive effect on many plants through recycling, such positive effects will be larger for the most nutrient limited species. Our model should then be seen as an approximation of such heterogeneities in the positive effects. Next to the extension to complex networks, another important issue would be to explicitly consider spatial dynamics. Indeed, local niche construction processes may create spatial heterogeneities in nature. Obvious examples include ant gardens (Frederickson et al., 2005) in which whole plant species communities are modified by the combination of trophic and non trophic effects of ant species, or large scale patterns of vegetation as in the termite-driven hypothesis for Namibian fairy circles (Juergens, 2013; Pringle & Tarnita, 2017).

